# The impact of dietary protein and carbohydrates on gene expression in a generalist insect herbivore

**DOI:** 10.1101/2020.10.30.361386

**Authors:** Carrie A. Deans, Greg Sword, Heiko Vogel, Spencer Behmer

## Abstract

Nutrition fuels all of the physiological processes that animals rely on for survival and reproduction. Of all the nutrients that are required, dietary protein (p) and carbohydrates (c) have a primary role. Insect herbivores are capable of detecting amino acid and sugar concentrations in plant tissue via chemoreception and regulate their intake of these two macronutrients to reach an optimal protein:carbohydrate, or p:c, ratio, termed an intake target. A multitude of studies have shown that the two nutritional factors that have the strongest impact on insect survival and performance are dietary p:c ratio and total macronutrient content, which is the proportion of the diet made up by p and c and a proxy for energy content. Variations in these two dietary traits have strong unique and interactive effects on many insect life history traits, yet the mechanisms that mediate these effects are not well understood. While many studies have documented the effect of host plant usage on gene expression, differences in plant secondary compounds between plant species and tissue types have confounded efforts to understand nutritional contributions to transcriptional changes. This study is the first to document the transcriptional effects of dietary p:c ratio and total macronutrient content in a phytophagous insect, the polyphagous moth species *Helicoverpa zea*. Our results show that changes in dietary p:c ratio produced a rather limited transcriptional response, while total macronutrient content had more dramatic effects on gene expression. The invariable expression of many metabolic genes across diets also suggests that *H. zea* larvae employ a strategy of constitutive expression to deal with nutritional imbalances rather than diet-associated changes in expression. We also observed many similarities in the transcriptional response to diets that varied from the intake target diet in different ways (c-biased, p-biased, increased energy content). This indicates that similar mechanisms are used to deal with nutritional imbalances regardless of the direction of the imbalance, and further supports the importance of nutrient regulation.

**HIGHLIGHTS:**

- Variations in plant macronutrients can have strong impacts on herbivore fitness
- Despite a wealth of studies documenting the physiological effects of macronutrient nutrition, underlying mechanisms are still ambiguous
- Diet protein-to-carbohydrate ratio had an unexpectedly small impact on overall transcription, while total macronutrient content had a stronger effect
- The transcriptional response to dietary variations away from an optimal diet was similar across diets that varied in different ways (carbohydrate-biased, protein-biased, more concentrated)
- Maintaining consistent consumption and constitutive expression of digestive enzymes across diets that varied in macronutrient profiles led to compensation for the most limiting dietary macronutrient

## INTRODUCTION

The ability of living things to obtain the nutrients they need from their environment has strong impacts on individual survival and performance but also has important implications for higher order processes, such as population dynamics, ecological interactions, and evolution. Despite the diversity that exists across the animal kingdom in size, shape, and life-history, all living things share a common requirement for dietary protein and carbohydrates. These macronutrients are required in the greatest amounts and consequently are most likely to be limiting in nature. To contend with this, animals have evolved sophisticated behavioral and physiological processes for detecting, regulating, and utilizing dietary protein and carbohydrates. Determining how these processes work together to mitigate the effects of nutritional variability is integral for understanding the mechanisms of nutritional plasticity and elucidating the connection between nutrition and animal fitness.

Insect herbivores routinely deal with considerable nutritional variability. Plant nutrient content is notoriously dynamic, varying both spatially across different plant families and individuals, but also temporally across seasonal and ontogenic changes. Leaf protein content alone can vary between 1-40% of dry mass (Bernays and Chapman, 1994), however, reproductive tissues typically exceed that of foliar tissues (Slansky and Scriber, 1984; Bernays and Chapman, 1994; Deans et al., 2016, 2018). Plant protein content also tends to decrease as tissues age throughout the growing season and can be increased significantly through fertilization and other environmental effects (Slansky and Scriber, 1984; Bernays and Chapman, 1994; Lenhart et al., 2015; Deans et al., 2016). Plant carbohydrate content is also highly variable, particularly between plant species that utilize different photosynthetic pathways, seasonal changes in light, temperature, and precipitation, and plant ontogeny (Bernays and Chapman, 1994; Lenhart et al., 2015; Deans et al., 2016, 2018).

Despite this nutritional heterogeneity, virtually all insect herbivores actively regulate their intake of dietary protein and carbohydrates to achieve a specific ratio, termed an intake target. The intake target represents the balance of dietary protein and carbohydrate intake that supports optimal performance and maximizes fitness (Simpson and Raubenheimer, 1993; Behmer, 2009). When insect herbivores are not able to reach their intake target, it can lead to reductions in fitness; however, this isn’t always the case. Many caterpillar species, particularly generalists, exhibit impressive nutritional plasticity and are able to maintain performance when feeding on diets that diverge considerably from their intake target. Deans et al. (2015) reared *Helicoverpa zea* larvae on artificial diets that mimicked the range of protein-to-carbohydrate (p:c) ratios and total macronutrient concentrations found in cotton (a common resource for *H. zea* larvae) and found remarkably few differences in developmental time, pupal mass, and growth rate, despite a p:c range of 0.4 to 2.5 and an over 3-fold difference in total macronutrient concentration. A similar result was seen for pupation success, developmental time, and pupal mass in the control treatments of Deans et al. (2017), which tested similar diets. Although several other caterpillar studies have documented significant effects of diet p:c on larval performance, most of these effects were only seen on the most extreme, and largely unnatural, diets. Performance across diet treatments that correspond to more field-relevant p:c ratios, like those tested by Deans et al. (2015, 2017), showed few significant differences and/or differences with limited biological significance (Lee et al., 2004; Thompson and Redak, 2005; Despland and Noseworthy, 2006; Lee et al., 2006; Merkx-Jacques et al., 2008).

Despite a wealth of data connecting dietary protein and carbohydrates to insect herbivore fitness, the mechanisms that mediate these effects are still ambiguous. Transcriptomic analyses offer the means to explore diet-mediated effects on gene expression and the opportunity to identify molecular and physiological mechanisms associated with nutritional plasticity. The affordability of next-generation sequencing and advancements in *de novo* techniques have made gene expression studies more feasible for non-model organisms; however, of the handful of studies that have measured gene expression or gene products across insect diets, there has been little consistency among the dietary factors tested. Several studies have focused on gene expression across different host plants or undefined artificial diets (Chougule et al., 2005; Osborn, 2011; Gog, 2012; Celorio-Mancera et al., 2012; Vogel et al. 2014) without documenting the macronutrient differences between them, and of these, the focus has largely been on the effects of plant secondary compounds (Govind et al., 2010; Oppert et al., 2010; Celorio-Mancera et al., 2011; Alon et al. 2012; Gog, 2012; Vogel et al., 2014) or protease-inhibitors (Moon et al., 2004; Chougule et al., 2005; Erlandson et al., 2010) rather than nutritional components. Our study is the first to explicitly measure transcriptional effects in response to quantifiable changes in diet macronutrients (p:c ratio and total macronutrient content) in a phytophagous insect.

In this study, we utilized RNA-seq to measure whole-body gene expression in *Helicoverpa zea* (also referred to as cotton bollworm, corn earworm, etc.) larvae fed on artificial diets that varied in both p:c ratio and total macronutrient content in order to identify gene functions and pathways that respond to differences in macronutrient profiles. Diet treatments consisted of artificial diets that were altered to match the p:c ratio and total macronutrient concentration of different plant tissues in two common hosts of *H. zea*-cotton (*Gossypium hirsutum*) and sweet corn (*Zea mays* L.) (Deans et al., 2016, 2018), as well as the self-selected intake target reported for *H. zea* (Deans et al., 2015). Table 1 shows the name of each diet treatment and its association to relevant plant tissues. Ultimately, we tested three different p:c ratios-a protein-biased ratio (PB-42), a carbohydrate-biased ratio (CB-42), and a ratio that approximated the self-selected intake target reported for *H. zea* (IT-42)-all at a total macronutrient concentration of 42%, which is a moderate macronutrient concentration for agricultural crops (Deans et al., 2016, 2018). We also tested a fourth diet that contained the same p:c ratio as the intake target but at a higher total macronutrient concentration of 68%, which is more indicative of reproductive tissues like cotton seed (IT-68). The diet treatments tested in this study also directly correspond to the control treatments tested in Deans et al. (2017), where larval performance was measured across these diets. Taken together, these studies allow for comparisons to be made between diet-specific gene expression profiles and larval performance.

**Table 1.** Treatment information. The associations between the treatment artificial diets, their p:c ratio and total macronutrient concentration, and the host plant tissues that correspond to each macronutrient profile (Deans et al., 2015, 2017, 2018).

| Treatment | P:C Ratio | Total Macro Content | Reference |
| --- | --- | --- | --- |
| protein-biased<br>(PB-42) | 30:12 (2.4) | 42% | cotton leaves |
| intake target<br>(IT-42) | 24:18 (1.3) | 42% | cotton squares (pre-flowers), self-selected IT |
| carb-biased<br>(CB-42) | 12:30 (0.4) | 42% | sweet corn kernels |
| intake target<br>(IT-68) | 39:29 (1.3) | 68% | developing cotton seed, self-selected IT |

Given that *H. zea* larvae have demonstrated an ability to maintain larval performance across diets that vary in p:c ratio and total macronutrient concentration (Deans et al., 2015, 2017), we hypothesized that considerable transcriptional activity would be required to mitigate nutritional variability. As such, we expected to see a high number of genes being differentially-expressed across both diet p:c comparisons and macronutrient concentrations. In particular, we expected to see some compensation in the expression of digestive enzymes, with the expectation that genes encoding proteases would likely be up-regulated in larvae fed the CB-42 diet and those encoding glycosidases would be up-regulated in larvae fed the PB-42 diet compared to the IT-42. Additionally, we anticipated that genes involved in energy metabolism would be up-regulated in the IT-68 compared to the IT-42. Because plant macronutrients are spatially and temporally variable and nutrition has strong impacts on insect performance, understanding the molecular mechanisms that mediate nutritional effects will be useful for predicting population dynamics in response to environmental variability, such as climate change, for developing effective pest control strategies, and for understanding insect evolution and host plant adaptation.

## METHODS

### Insects

Caterpillar eggs (*Helicoverpa zea*) were obtained from Benzon Research (Carlisle, PA). Neonates were hatched in the laboratory and transferred to individual cells in plastic trays using a fine-tipped paint brush. Throughout the experiment larvae were housed in a growth chamber Model I-66VL; Percival Scientific, Perry, IA, USA) set to 25°C with a 14:10 L:D cycle.

### Treatments

Neonates (n=10) were assigned to one of four diet treatments, previously described as the control treatments in Deans et al. (2017). Table 1 shows the name of each diet treatment, its p:c ratio, total macronutrient concentration, and the cotton tissue it corresponds to, while Figure 1 shows the location of each treatment in nutrient space. These diets were selected because they represent a range of p:c ratios and total macronutrient concentrations that are field-relevant, in that they correspond the macronutrient profiles of different cotton and sweet corn tissues as outlined in Deans et al. (2016) and Deans et al. (2018). Because sweet corn and cotton are common hosts for *H. zea*, these diets simulate the nutritional variability encountered by larvae in the field. Treatments consisted of a three p:c ratios with a total macronutrient concentration of 42%, which is indicative of vegetative tissues: a carbohydrate-biased diet (CB-42) mimicking sweet corn kernels, a diet that approximated the reported intake target for *H. zea* (IT-42) (Deans et al. 2015) and reflected the nutrient content of immature cotton flowers, or squares, as well as a protein-biased diet (PB-IT) that represented the nutrient content of mature leaves. The intake target ratio was also tested at a higher total macronutrient concentration of 68% (IT-68), which is indicative of reproductive tissues and mimicked the nutrient content of developing cotton seeds, a preferred resource for *H. zea* larvae. Larvae were reared on their respective diets during their entire larval development and fed *ad libitum*. Fresh diet was provided at a minimum of every four days.

**Fig. 1.**
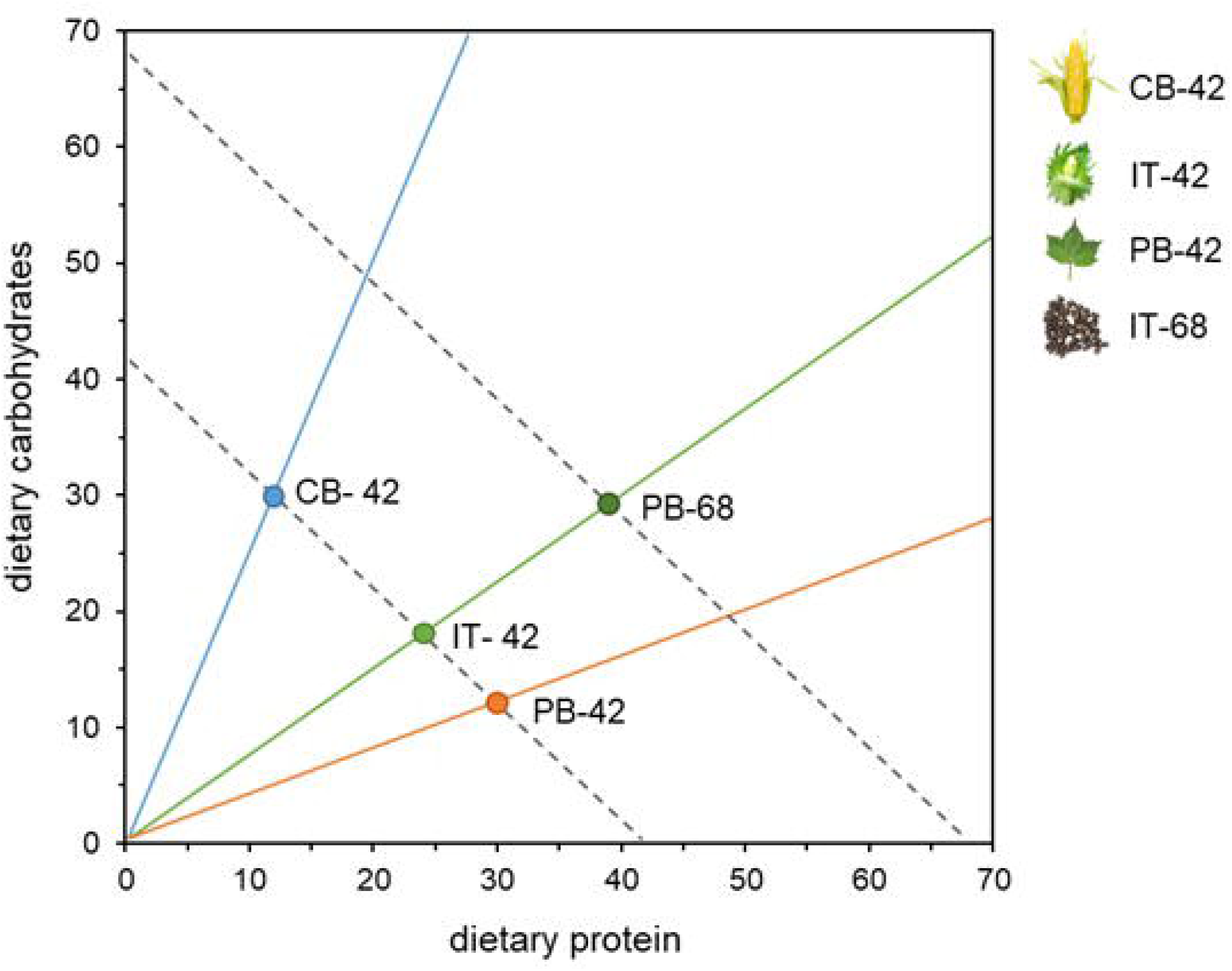
Treatments in nutrient space. Bi-coordinate plot showing the location of each diet treatment in nutrient space delineated by dietary protein and carbohydrate content. The legend also shows the cotton tissue that corresponds to each treatment macronutrient profile based on Deans et al. (2016). Iso-caloric lines (dotted) are shown for diets with 42% and 68% total macronutrient content.

### Protocol

Twelve hours after molting to the 4^th^ instar, larvae were dipped in liquid nitrogen and stored individually at −80°C. Frozen larvae were crushed with a disposable pestle in a 1.5mL microcentrifuge tube and mixed with 1 mL of TRIzol reagent. Due to the exploratory nature of this study, whole-body samples were used; however, midgut tissue likely accounted for the majority of larval biomass. RNA extractions were then done on a 50% dilution of this mixture (0.5 mL of mixture added to 0.5 mL of TRIzol reagent). Extractions were done using Direct-zol™ RNA MiniPrep columns (Zymo Research) with 10uL elutions. Extractions of four replicates from each treatment (16 samples total) were sent to the Texas A&M University’s Genomics and Bioinformatic Service for quality analysis and sequencing. Sixteen strand-specific libraries were prepared using Illumina TruSeq library prep and 150bp paired-end reads were sequenced using an Illumina HiSeq2500. Library fragment size selection was kept small (75bp) to reduce the exclusion short-sequence immunity genes.

### Bioinformatics

Tophat (Trapnell et al., 2013) was used to align processed reads to an early version of the *H. zea* genome (provided by Tom Walsh, CSIRO). The complete genome is now available on the NCBI website (accession #: GCA_002150865.1) (Pearce et al., 2017). FactoMineR was used in R to create a PCA plot to show individual variation. A differential gene expression analysis was done using edgeR within BLAST2GO (GLM likelihood ratio test with an FDR threshold of 0.05 and a log fold-change threshold of 1) (Götz et al., 2008; Robinson et al., 2010). Contrasts were made between the intake target diet and the other three diets, resulting in three different comparisons: CB-42 vs IT-42, PB-42 vs IT-42, and IT-68 vs IT-42. The CB-42 vs IT-42and the PB-42 vs IT-42 contrasts identified genes related to changes in dietary p:c ratio, while the IT-68 vs IT-42 contrast identified genes related to changes in total macronutrient content. Differentially-expressed (DE) gene sequences were BLASTed (blastp-fast) against the non-redundant (nr) ncbi database and were cross-referenced against the InterPro database using the public EMBL-EMI InterPro web-service. The BLAST results were then mapped using data from the Gene Ontology Association and UniProt database. All DE, mapping, and annotation analyses were done using BLAST2GO (Kanehisa et al., 2000).

For each diet contrast, up- and down-regulated DE genes were annotated with gene ontology (GO) terms using BLAST2GO. Gene ontology was compared across contrasts in two ways. First, a Chi-square test was used to determine whether the number of up- and down-regulated genes within each GO category differed between contrasts. Second, an enrichment analysis (Fisher’s exact test) was performed in BLAST2GO to identify over-represented GO terms for each contrast. These are GO categories that include a disproportionately higher number of DE genes in a particular contrast as compared to the total number of genes in that GO category present in the entire genome. Differentially-expressed genes sets were also manually annotated based on gene name and GO to identify genes involved in protein digestion (trypsin, chymotrypsin, aminopeptidase, and carboxypeptidase), carbohydrate digestion (amylase, alpha-N-acetylgalactosaminidase, trehalase, lactase, and maltase), and lipid digestion (lipase), as well as macronutrient transport. A Gene Set Enrichment Analysis (GSEA) was also done using GSEA (GSEA 3.0, Broad Institute) to determine whether these categories of genes were correlated with specific diet treatments. Gene sets were manually curated by searching for genes within the *H. zea* genome that contained the relevant names and/or GO terminology. In GSEA, the expression of each gene in a gene set is compared across contrasts to determine if the gene set in question is significantly correlated with a specific treatment. Gene sets are comprised of genes with specific putative gene functions, whether they have been shown to be DE or not. In this way, GSEA looks at cumulative expression across specific gene categories to determine if that function is associated with a particular treatment, even if the expression of individual genes within those categories are not DE.

To determine the connection between the diet treatments and the expression of genes related to specific metabolic pathways, we utilized consumption data from Deans et al. (2017) to calculate mass-specific protein and carbohydrate consumption across diets. We then combined consumption data with the total expression of protease (trypsin, chymotrypsin, aminopeptidase, and carboxypeptidase) and glycosidase (amylase, alpha-N-acetylgalactosaminidase, trehalase, lactase, and maltase) genes to estimate the total expression (in transcripts-per-million, or TPM) of protease or glycosidase genes per average protein or carbohydrate consumption measured on a mass-specific basis (mg p/mg insect/day). This allowed us to account for protein and carbohydrate consumption when assessing differences in the transcriptional response of specific digestive enzymes We used an ANOVA (SPSS Statistics 24 Inc., Chicago, IL, USA) to detect significant differences in mass-specific protein and carbohydrate consumption across diets, as well as differences in consumption-based expressional changes.

## RESULTS

### Summary Statistics

Across all samples we obtained 320,277,088 total reads, with an average of 20,017,318 reads per sample. About 90% of our reads successfully mapped to our reference genome, and over 80% of our read pairs showed concordant alignments between pairs. We were able to obtain BLAST hits for almost all of the differentially-expressed (DE) sequences (98%) and obtain annotation information for about 70% (SI Fig. 1). SI Fig. 2 shows the individual variation between replicates within each diet treatment. Clustering was apparent, with the PB-42 exhibiting the tightest clustering among replicates and the IT-42 and IT-68 having a greater spread.

**Fig. 2.**
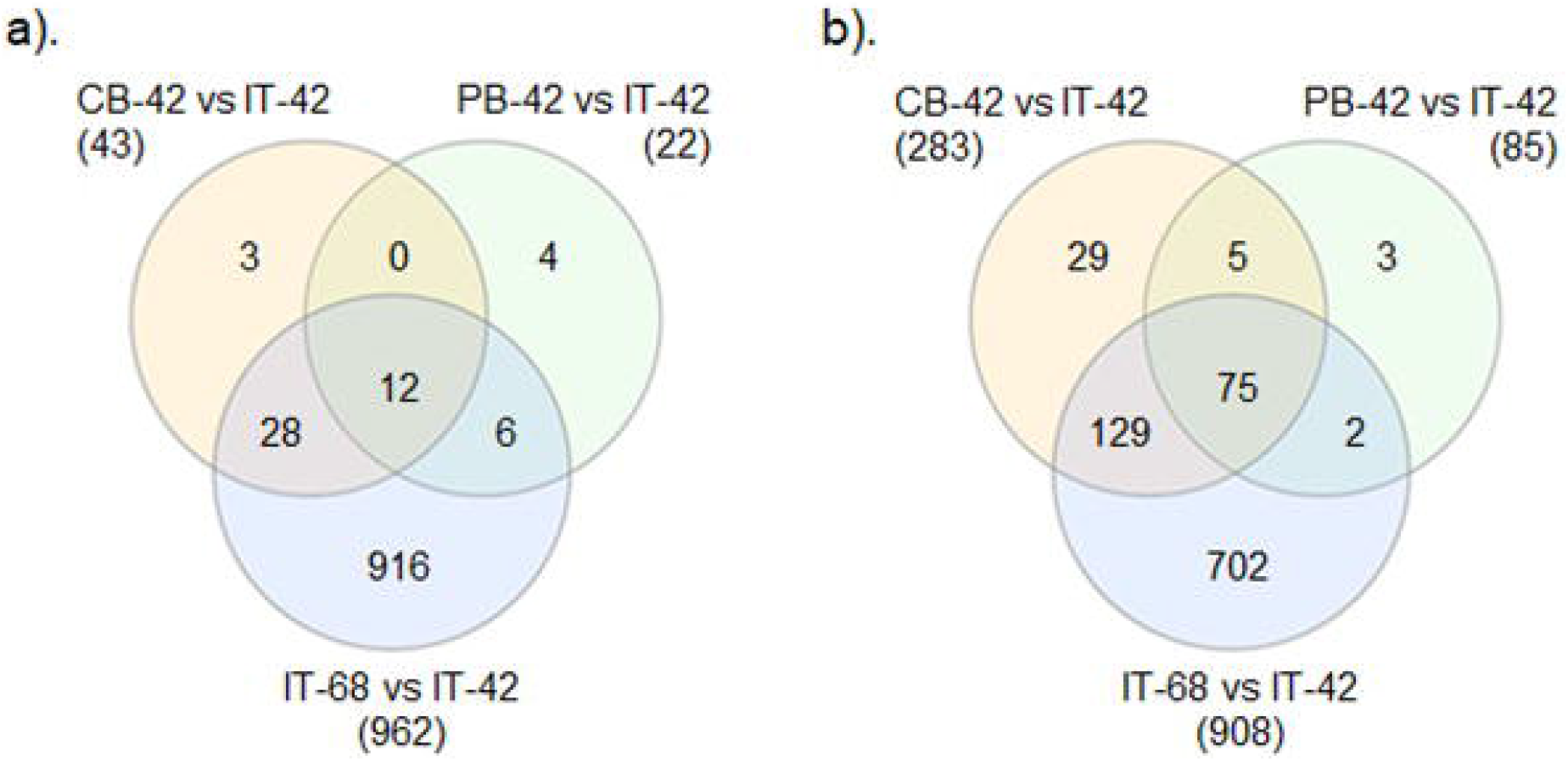
Differential gene expression. Venn diagram showing the number of differentially-expressed genes found for each contrast. (a) Up-regulated and (b) down-regulated genes.

### Diet Effects

To determine the transcriptional effects of diet p:c ratio and total macronutrient concentration, contrasts were made between the self-selected and presumably optimal IT-42 diet and each of the other three diets. Comparisons between the CB-42 and IT-42 diets and between the PB-42 and IT-42 diets allowed us to assess the effects of variations in dietary p:c ratio, while comparisons between the concentrated IT-68 diet and the less concentrated IT-42 diet allowed us to assess the effects of variation in total macronutrient concentration.

#### CB-42 vs IT-42

Feeding on the carbohydrate-biased CB-42 diet resulted in the differential regulation of 281 genes (Figure 2a). The majority of these genes, about 85%, were down-regulated while 15% were up-regulated. Up-regulated DE genes represented a total of 39 GO terms. Of those terms, membrane, catalytic activity, transporter activity, localization, and biological regulation were significantly associated with the up-regulated genes (Figure 3). Down-regulated DE genes represented a total of 142 GO terms. The GO terms significantly associated the genes down-regulated in the CB-42 treatment included many terms related to protein-intensive processes, such as protein-containing complex, antioxidant activity, detoxification, immune system process, and reproduction (Figure 4). The Enrichment Analysis (EA) showed that processes related to carbohydrate metabolism and general transport were enriched among up-regulated genes, while those related to oxidation-reduction, chitin metabolism, and binding were overrepresented among the down-regulated DE genes (Figure 5a-b).

**Fig. 3.**
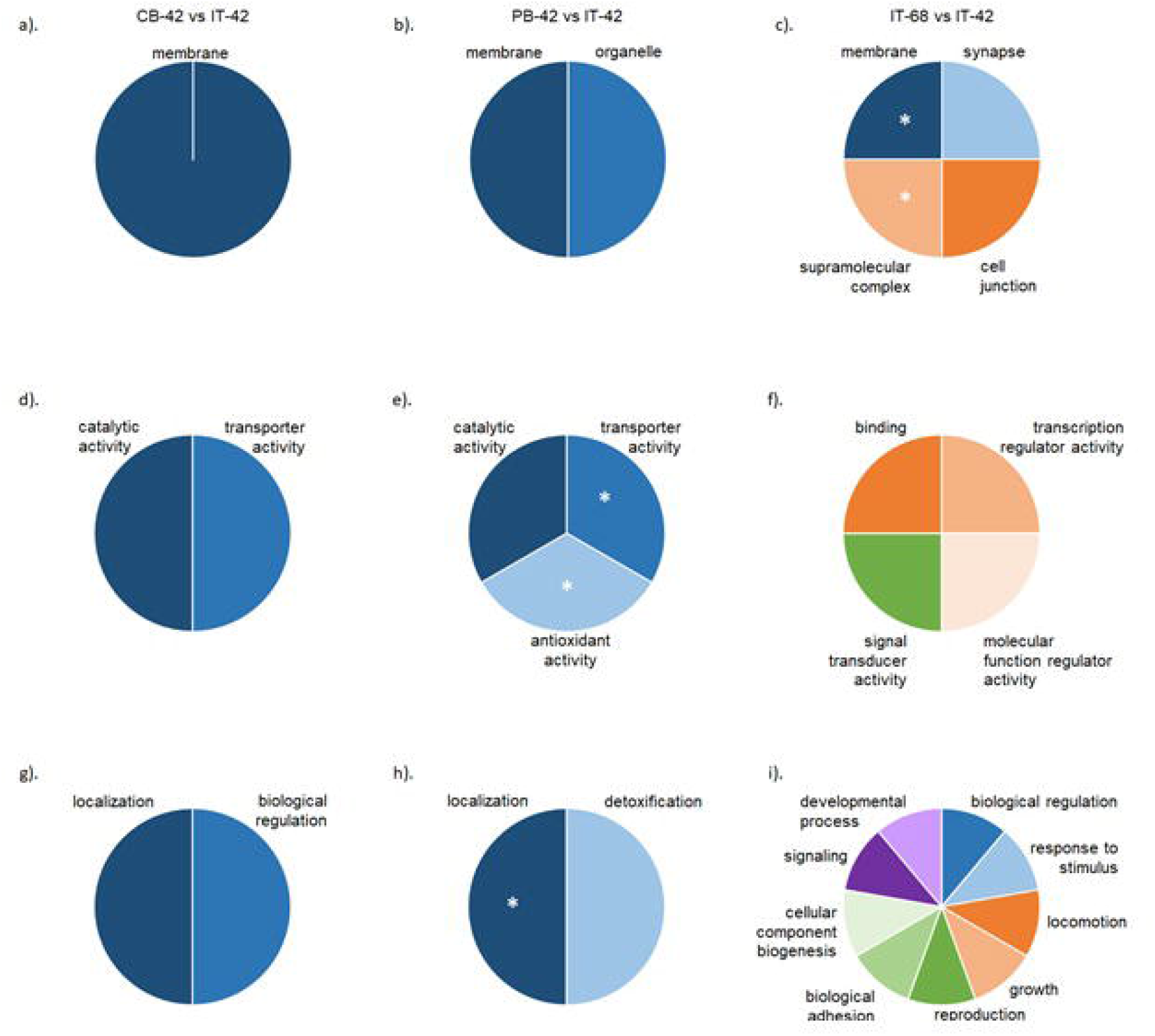
Gene ontology for up-regulated genes. Pie charts showing the proportion of DE genes associated with each significantly up-regulated GO terms for each diet. Each row shows terms associated with cellular components (a-cl, molecular function (d-f), and biological processes (g-i). GO terms marked with an * identify terms that were uniquely up-regulated while the other terms were found to have a significantly higher number of genes up-regulated than down-regulated as indicated by a Chi-square test (P ≤ 0.05).

**Fig. 4.**
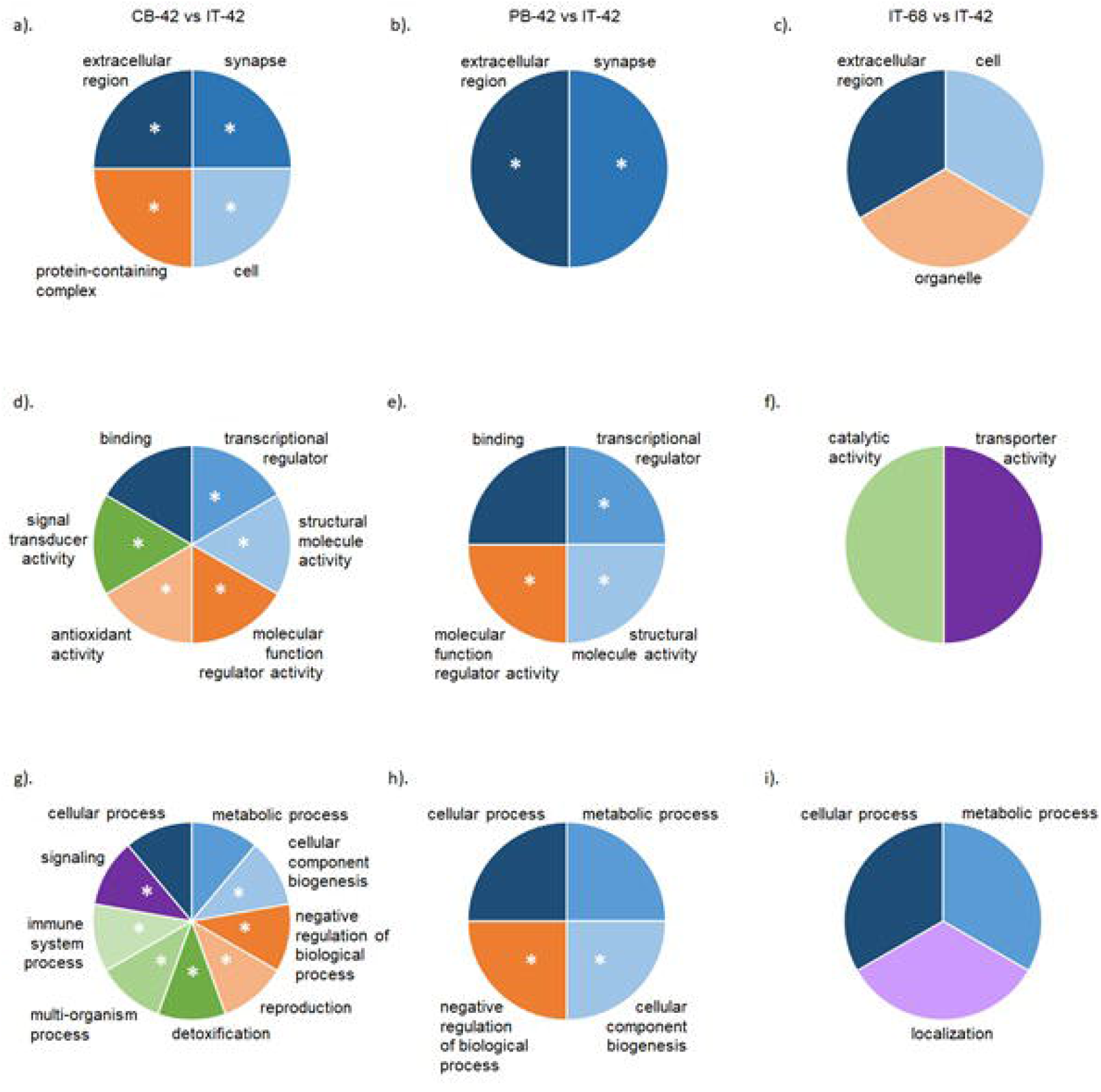
Gene ontology for down-regulated genes. Pie charts showing the proportion of DE genes associated with each significantly down-regulated GO terms for each diet. Each row shows terms associated with cellular components (a-c), molecular function (d-f), and biological processes (g-i). GO terms marked with an * identify terms that were uniquely down-regulated while the other terms were found to have a significantly higher number of genes down-regulated than up-regulated as indicated by a Chi-square test (P ≤ 0.05).

**Fig. 5.**
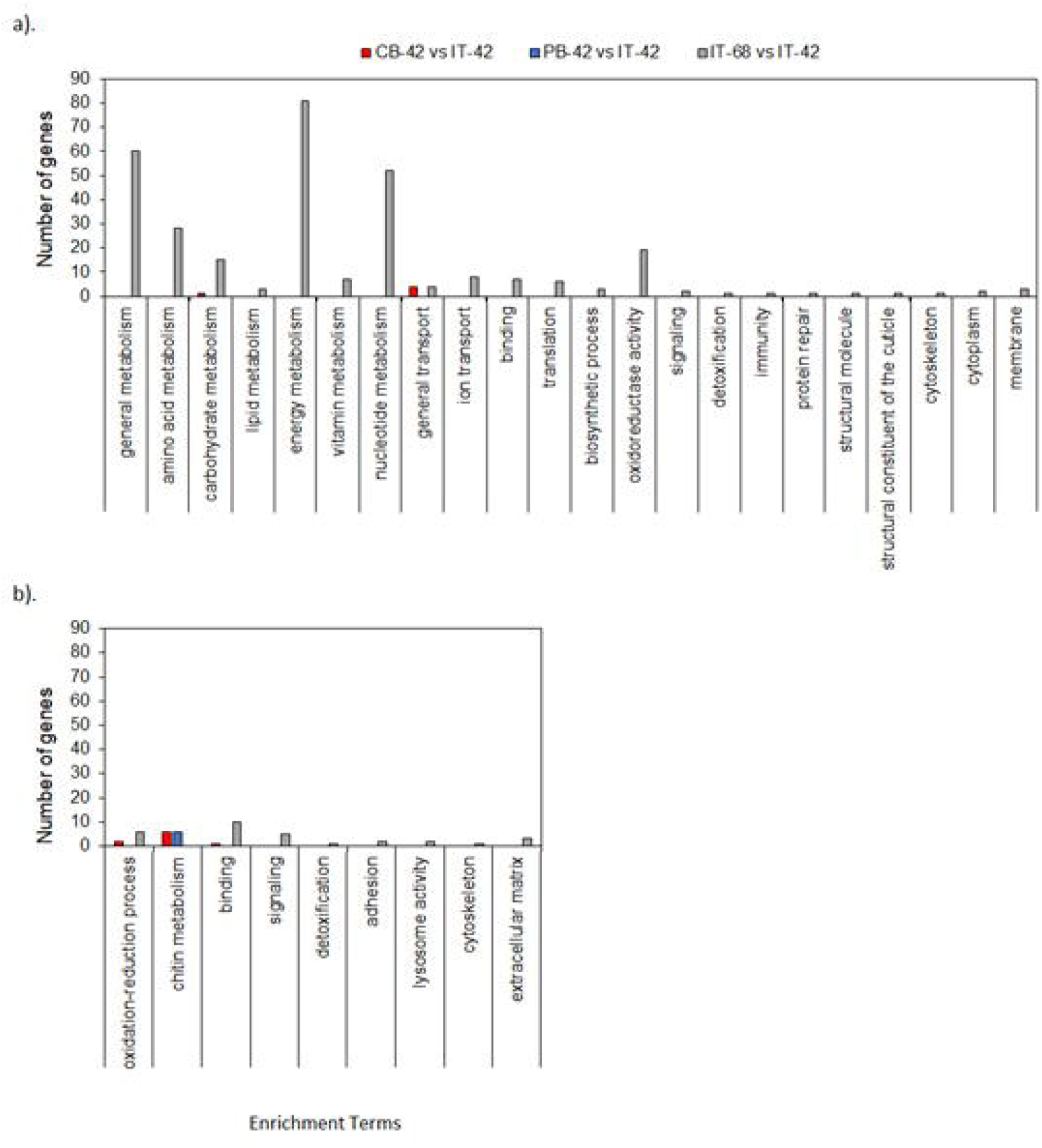
Enrichment analysis results. Bar graphs showing the GO terms that showed significant enrichment (FDR ≤ 0.05) across diets and the number of genes associated with each term. (a) Terms enriched among up-regulated DE genes, (b) terms enriched among down-regulated DE genes.

Manual annotation identified few up-regulated genes related to digestive enzymes; however, 14 protease genes were found to be down-regulated in CB-42 treatment (Figure 6). Despite a rather high number of protease genes being down-regulated, the GSEA did find a significant correlation between the expression of aminopeptidase genes and the CB-42 treatment (Table 2). Table 2 shows that genes related to energy metabolism were also significantly correlated to the CB-42 treatment.

**Table 2.**
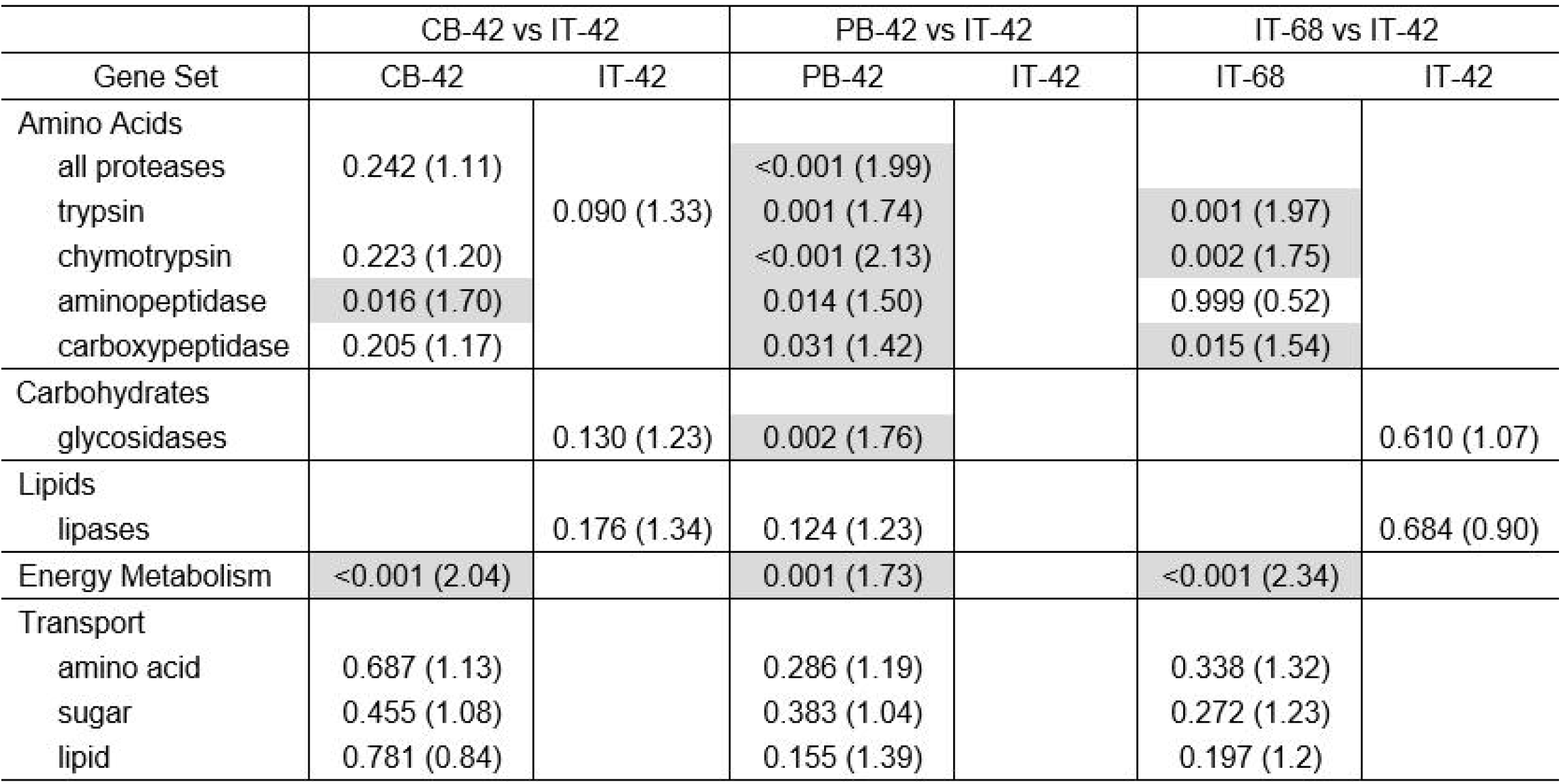
GSEA results. The FDR values, with absolute value of the normalized enrichment scores (NES) in parentheses, for each gene set. Darkened cells indicate the genes sets that were significantly enriched (FDR ≤ 0.05).

**Fig. 6.**
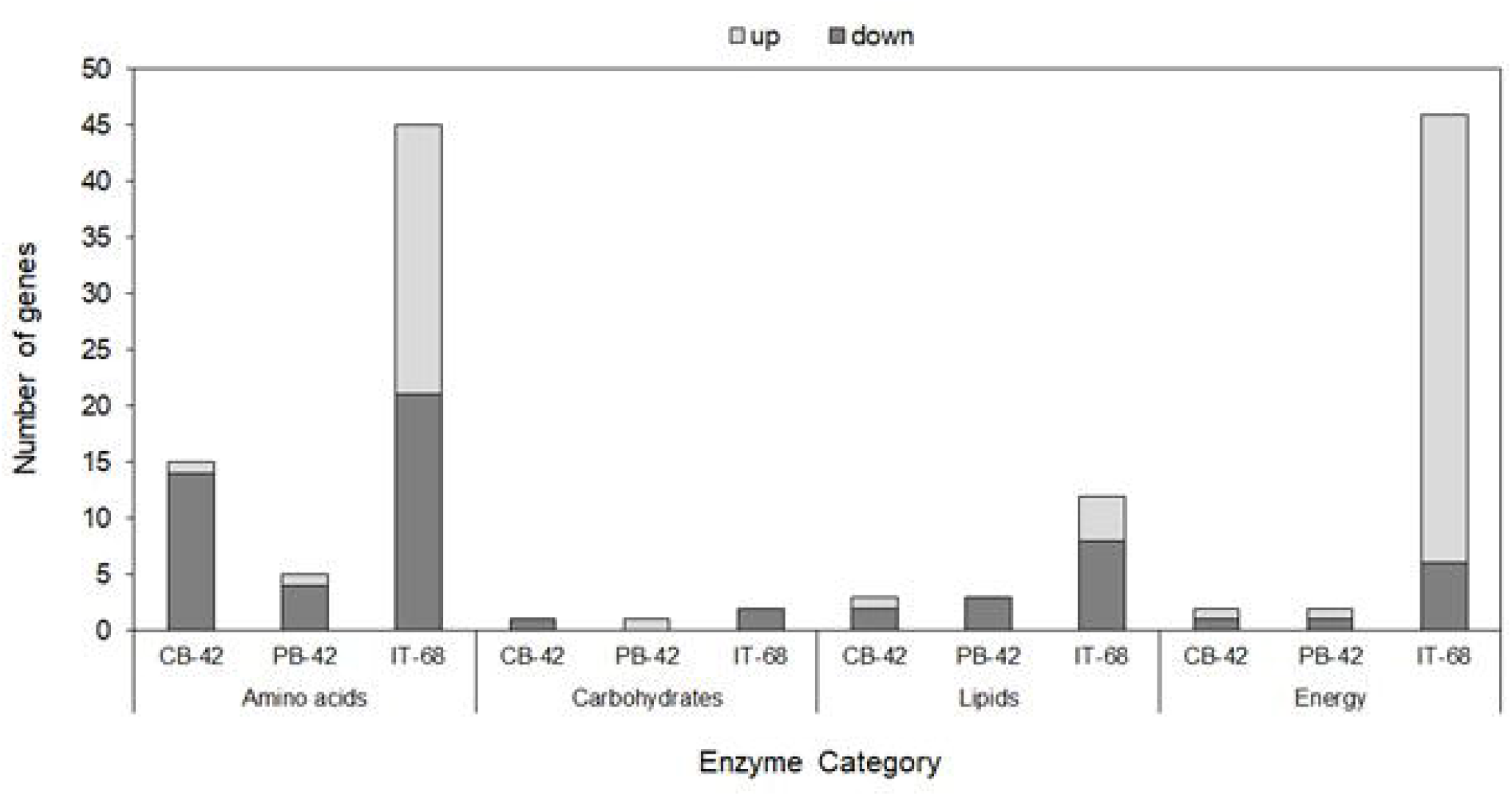
Enzyme specific differentially-expressed genes. Bar graphs showing the number of up- and down-regulated differentially-expressed genes associated with specific digestive enzymes or metabolic pathways for each diet contrast. Genes were manually annotated based on gene name and GO terminology.

#### PB-42 vs IT-42

Feeding on the protein-biased PB-42 diet impacted the regulation of 107 genes, with about 80% of DE genes showing down-regulation and 20% up-regulation (Figure 2a-b). The up-regulated DE genes corresponded to 55 GO terms. Terms significantly associated with up-regulation were related to the membrane, organelle, catalytic activity, transporter activity, localization, and the protein-intensive processes of antioxidant activity and detoxification (Figure 3). Down-regulated DE genes contained 87 GO terms, with significant terms relating to: the extracellular matrix, synapse, binding, cellular and metabolic processes, transcription, structural molecule activity, molecular function and biological regulation, and cellular component biogenesis (Figure 4). Despite these GO term associations, the EA found no terms enriched among the up-regulated DE genes and only one term, chitin metabolism, to be enriched among the down-regulated DE genes in the PB-42 treatment (Figure 5a-b).

The PB-42 diet showed few DE genes associated with digestive enzymes. Figure 6 shows the presence of only 1 up-regulated gene related to amino acid digestion and 1 gene related to carbohydrate digestion. Four genes associated with amino acid digestion and 3 genes related to lipid digestion were found among down-regulated DE genes for the PB-42 treatment. Despite finding a limited number of DE genes encoding digestive enzymes, the GSEA showed that expression levels among gene sets related to various proteases were significantly correlated with feeding on the PB-42 diet (Table 2). Table 2 also shows that glycosidase expression and the expression of genes related to energy metabolism were also significantly correlated to the PB-42 diet compared to the IT diet.

#### IT-68 vs IT-42

Compared to the p:c ratio contrasts, feeding on a diet with a high total macronutrient content had a greater impact on overall transcription. Simply changing the energy content of the IT diet from 42% to 68% resulted in the regulation of 1,870 genes. The number of up-regulated versus down-regulated genes were similar, with 51% of genes up-regulated and 48% down-regulated (Figure 2a-b). The up-regulated genes contained 248 GO terms. Of those, the terms significantly up-regulated were related to the synapse, supramolecular complex, cell junction, binding, transcriptional regulator activity, molecular function regulator, signal transducer activity, response to stimulus, locomotion, growth, reproduction, biological adhesion, cellular component biogenesis, signaling, and developmental process (Figure 3). Down-regulated genes corresponded to 198 GO terms, with significantly down-regulated terms including extracellular region, cell, organelle, catalytic activity, transporter activity, cellular and metabolic processes, and localization (Figure 4). The EA indicated that 22 GO term were enriched among up-regulated genes in the IT-68 treatment, including general metabolism, amino acid, carbohydrate, and energy metabolism, and to a lesser extent lipid metabolism (Figure 5a). Nine terms were enriched among down-regulated genes, including oxidation-reduction process and signaling (Figure 5b).

Figure 6 shows that manual annotation found 24 genes related to amino acid digestion and 4 genes related to lipid digestion among the up-regulated genes in the IT-68 treatment, while 40 genes related to amino acid digestion and 8 genes related to lipid digestion were found among the down-regulated genes. Forty genes related to energy metabolism were up-regulated while only 6 were down-regulated. The GSEA showed a strong correlation between the IT-68 treatment and all protease gene sets, except aminopeptidases, as well as energy metabolism genes (Table 2).

### Diet Comparisons

Figure 2a shows that despite showing a total of 65 up-regulated DE genes across both treatments, no genes were shared among the CB-42 and PB-42 diet contrasts. Alternatively, the CB-42 diet shared 65% of its up-regulated genes (28 genes) with the IT-68 diet, which contained a similar percentage of dietary carbohydrates. These shared genes were broadly associated with energy metabolism, sugar transport, iron binding, and detoxification. The PB-42 diet shared only 27% of its up-regulated genes (6 genes) with the IT-68 diet, and these included genes related to energy metabolism, protein binding, cation transport, oxidation-reduction activity, and transferase activity (Table 3). Twelve genes were shared by all three diet contrasts and represent genes that are generally associated with feeding on a diet that diverges from the IT diet, either in p:c ratio or total macronutrient content. These genes were associated with ion, folate and amino acid transport, lipid metabolism, calcium sequestration/release, and a salivary metalloenzyme (Table 4). This left 7%, 18%, and 95% of up-regulated DE genes being unique to the CB-42, PB-42, and IT-68 diets, respectively.

**Table 3.** Consumption data from Deans et al. (2017). Mean mass-specific protein and carbohydrate consumption (± SE) for each diet (N). Mass-specific p (ANOVA: F3,72=92.82, P<0.0001) and c consumption (ANOVA; F3,72=86.95, P<0.0001) were significantly different. Different letters indicate significant post hoc differences between diets (P ≤ 0.05).

| Diet | Protein Cons. (mg/mg/day) | Carbohydrate Cons. (mg/mg/day) |
| --- | --- | --- |
| p12:c30 (CB-42) (16) | 0.0163 ± 0.00061 (a) | 0.0407 ± 0.00152 (a) |
| p24:c18 (IT-42) (26) | 0.0325 ± 0.00138 (b) | 0.0244 ± 0.00103 (b) |
| p30:c12 (PB-42) (17) | 0.0432 ± 0.00152 (c) | 0.0173 ± 0.00061 (c) |
| p39:c29 (IT-68) (17) | 0.0559 ± 0.00331 (d) | 0.0416 ± 0.00246 (a) |

**Table 4.** Statistics for protease and glycosidase expressional differences. ANOVA results for total protease and glycosidase gene expression (TPM) and consumption-specific protease and glycosidase gene expression (TPM/mass-specific p or c consumption (mg/mg/day)) across diets. Bolded values indicate statistical significance and *P* ≤ 0.05.

| Digestive Enzymes | Genes | Df | F Statistic | P-value |
| --- | --- | --- | --- | --- |
| Protease | Total expression | 3, 12 | 2.55 | 0.105 |
|  | Consumption-specific expression | 3, 12 | 12.29 | **0.001** |
| Glycosidase | Total expression | 3, 12 | 5.32 | **0.015** |
|  | Consumption-specific expression | 3, 12 | 11.72 | **0.001** |

Figure 2b shows that the CB-42 and PB-42 diets shared 5 down-regulated DE genes, a mere 1.5% of the total number of DE genes. Of these, only two annotations were found, both relating to calcium binding and the cuticle. The CB-42 diet again shared a rather large proportion, 54%, of their down-regulated DE genes with the IT-68 diet. These genes were involved in energy metabolism, chitin metabolism, lipid and co-factor metabolism, cuticle formation, the cytoskeleton, immunity, membrane components, calcium binding and transport, copper and ion binding, sugar and iron transport, ubiquitination, signaling, transcription, hormone production, cell adhesion, the cell cycle, transferase and oxidase activity, neuronal development, detoxification, as well as several proteases. The PB-42 diet shared only 2 genes, or 2% of its total down-regulated DE genes, with the IT-68 diet. These genes were involved in membrane components and transferase activity (Table 3). In total, 75 genes were shared across all diets, and these included several genes involved in lipid metabolism, chitin metabolism, serine-type endopeptidase activity, and cuticle formation (Table 4). This resulted in 12% of genes being uniquely expressed in the CB-42, 3.5% in the PB-42, and 77% in the IT-68 diets. The large portion of uniquely-expressed DE genes in the IT-68 diet is largely due to the higher number of DE genes found in the contrast with this diet compared to the CB-42 or PB-42 diets.

#### Consumption-Specific Transcriptional Effects

In order to account for the effect that differences in protein and carbohydrate consumption may have had on transcription across diet treatments, we used consumption data from Deans et al. (2017) to recalculate expression of protease and glycosidase genes on the basis of mass-specific consumption. Deans et al. (2017) reported that diet had no significant effect on total consumption or mass-specific diet consumption for larvae on control diets (which correspond to the p:c and total macronutrient content tested in our diet treatments); however, mass-specific protein (ANOVA: *F_3,72_*=92.82, *P*<0.0001) and carbohydrate consumption (ANOVA; *F_3,72_*=86.95, *P*<0.0001) were significantly different due to the differences in the protein and carbohydrate content in the diets. Mass-specific protein consumption differed significantly between each diet, scaling with the proportion of dietary protein (Table 3). Mass-specific carbohydrate consumption also scaled significantly with dietary carbohydrate content (Table 3). The protein-biased diet matching our PB-42 diet showed the lowest mass-specific carbohydrate consumption, followed by the intake target diet (IT-42), while the carbohydrate-biased (CB-42) and concentrated intake target (IT-68) diets showed statistically similar consumption, likely due to the fact that they had a similar percentage of dietary carbohydrates (~30%).

Table 4 shows that the total expression of protease genes (trypsin, chymotrypsin, aminopeptidase, and carboxypeptidase) did not vary significantly across diet treatments (Figure 7a); however, after accounting for protein consumption, differences were apparent. Larvae from the CB-42 diet showed a significantly higher level of protease expression than the other three diets (Figure 7b). However, despite as much as a 15% difference in dietary protein content, protease expression was statistically similar across the IT-42, PB-42, and IT-68 diets.

**Fig. 7.**
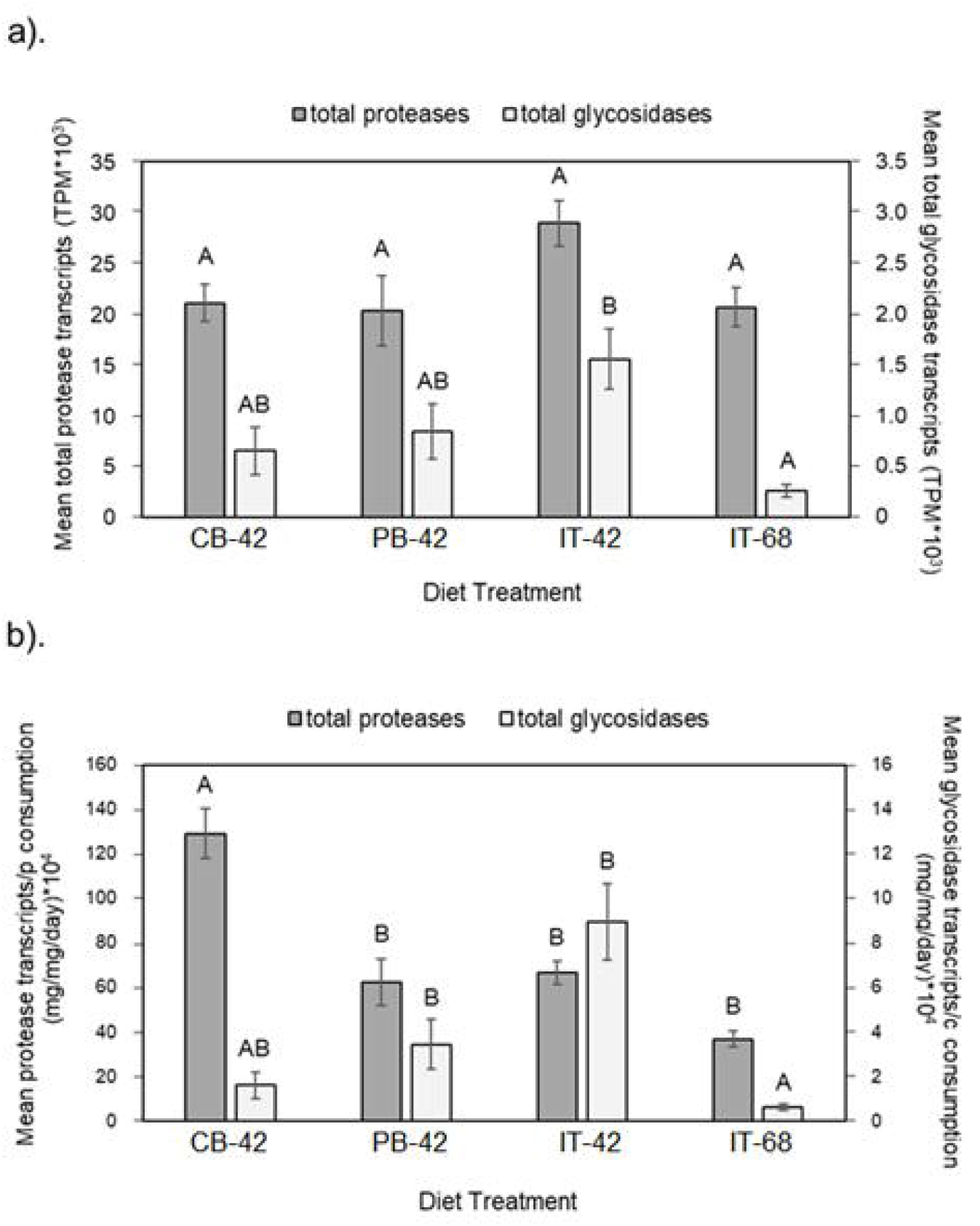
Total and consumption-specific expression of protease and glycosidase genes across diet. Bar charts showing the total number of average protease and glycosidase expression in larvae from each diet treatment (a) and protease and glycosidase expression adjusted for protein and carbohydrate consumption (b).

The total expression of glycosidase genes (amylase, alpha-N-acetylgalactosaminidase, trehalase, lactase, and maltase) only differed significantly between the PB-42 diet and the IT-68 diet, but all the 42% total macronutrient diets showed similar expression (Figure 7a). When carbohydrate consumption was taken into account, the concentrated IT-68 diet showed similar glycosidase expression to the carbohydrate-biased CB-42 diet but significantly lower expression compared to the IT-42and PB-42 diets (Figure 7b). Larvae fed on the PB-42 diet, which had the lowest mass-specific carbohydrate consumption, showed the highest expression of glycosidase transcripts per mg of carbohydrates consumed.

We also looked at the total expression of different types of proteases across diets and the total expression adjusted for protein consumption (Table 5). Although chymotrypsin was the only category to show significant differences in expression across diets (Figure 8a), when adjustments were made for protein consumption every protease category showed a significant difference in expression across diets (Figure 8b). The CB-42 diet showed significantly higher expression across all protease categories, except for aminopeptidases where expression was only higher than the IT-68 diet. Overall, expression was lowest for the IT-68 diet, although not significantly lower in all cases.

**Table 5.** ANOVA statistics for protease and glycosidase gene expression. Results for the effect of diet on the total expression and consumption-specific expression of different protease (all proteases, trypsin, chymotrypsin, aminopeptidase, and carboxypeptidase genes) and glycosidases (all glycosidases, including amylase, alpha-N-acetylgalactosaminidase, trehalase, lactase, and maltase genes) gene categories. Bolded values indicate statistical significance and P ≤ 0.05.

| Genes | Total Expression |  |  | Cons-Specific Expression |  |  |
| --- | --- | --- | --- | --- | --- | --- |
|  | Df | F Stat | P-value | Df | F Stat | P-value |
| all proteases | 3, 12 | 2.87 | 0.080 | 3, 12 | 22.414 | **<0.0001** |
| trypsin | 3, 12 | 2.42 | 0.117 | 3, 12 | 9.406 | **0.002** |
| chymotrypsin | 3, 12 | 5.61 | **0.012** | 3, 12 | 22.565 | **<0.0001** |
| aminopeptidase | 3, 12 | 1.702 | 0.219 | 3, 12 | 4.955 | **0.018** |
| carboxypeptidase | 3, 12 | 1.86 | 0.190 | 3, 12 | 19.603 | **<0.0001** |
| all glycosidases | 3, 12 | 5.477 | **0.013** | 3, 12 | 13.325 | **<0.0001** |
| α-amylase | 3, 12 | 8.830 | **0.002** | 3, 12 | 15.568 | **<0.0001** |
| α-trehalase | 3, 12 | 2.154 | 0.147 | 3, 12 | 9.188 | **0.002** |
| α-glucosidase | 3, 12 | 0.352 | 0.789 | 3, 12 | 3.028 | 0.071 |
| α-N-acetylgalactosaminidase | 3, 12 | 3.671 | **0.044** | 3, 12 | 7.324 | **0.005** |
| lactase | 3, 12 | 8.535 | **0.003** | 3, 12 | 9.268 | **0.002** |
| maltase | 3, 12 | 5.793 | **0.011** | 3, 12 | 7.878 | **0.004** |

**Figure 8.**
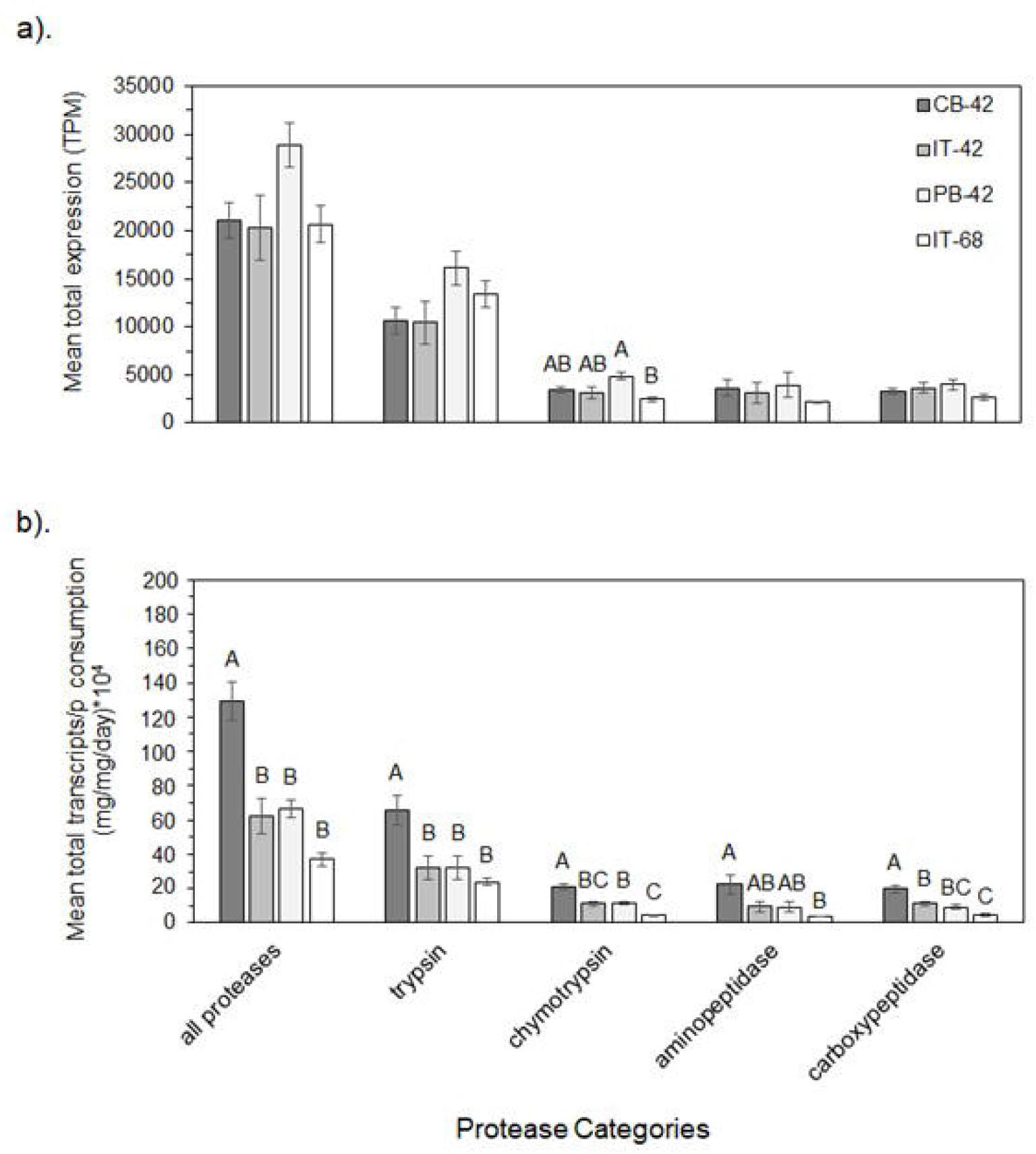
Expression of different proteases. Bar chart showing mean total expression of genes associated with different types of proteases (a) and the consumption-specific expression (b) for each diet treatment. Different letters indicate significant differences between diets within each category (P ≤ 0.05).

The expression of all glycosidases, α-amylases, α-N-acetylgalactosaminidases, and maltases were significantly different across diets (Table 5); however, more categories showed significant diet effects after accounting for carbohydrate consumption. Figure 9b shows that the protein-biased PB-42 diet consistently showed the highest expression across all categories, while both the CB-42 and IT-68 diets showed the lowest expression. Expression in the IT-42 diet was intermediate, but it wasn’t significantly different from that of the protein-biased diet for most glycosidase categories.

**Figure 9.**
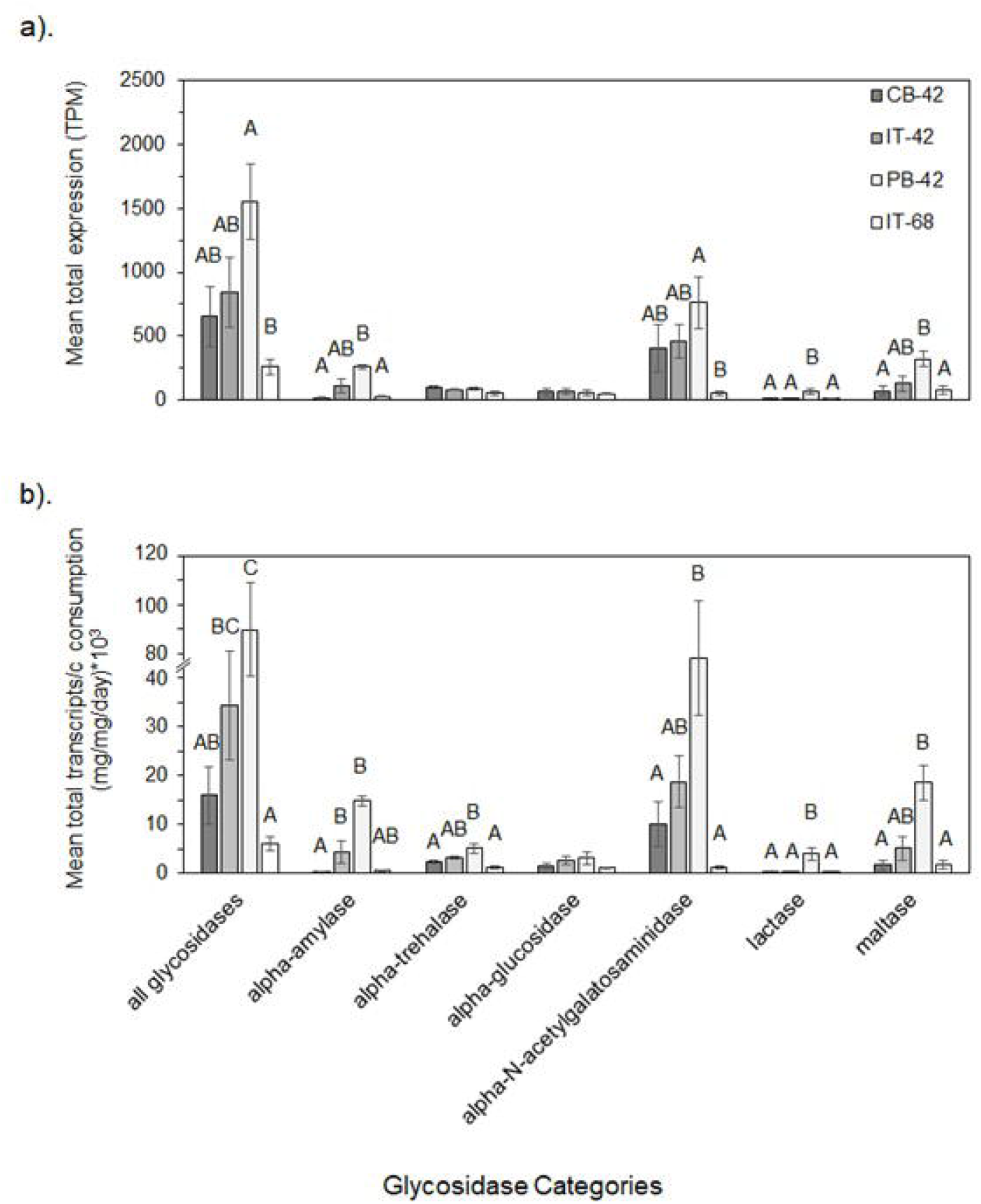
Expression of different glycosidases. Bar chart showing mean total expression of genes associated with different types of glycosidases (a) and the consumption-specific expression (b) for each diet treatment. Different letters indicate significant differences between diets within each category (P ≤ 0.05).

## DISCUSSION

This study is the first to explore how field-relevant variations in dietary p:c ratio and total macronutrient content impact gene expression in a herbivorous insect. Comparing the expression profiles of larvae fed a carbohydrate-biased or a protein-biased diet to those fed on a diet that approximated the intake target allowed us to examine the effect of p:c ratio on transcription. Additionally, comparing gene expression in larvae fed the intake target diet to those fed on a more concentrated diet with the same p:c ratio, allowed us to examine the impact of total macronutrient content on transcription. Because Deans et al. (2017) observed few differences in *H. zea* larval performance across control diets, we expected that maintaining performance across nutritional variability would require significant changes at the transcriptional level. As such, we hypothesized that the expression of metabolic genes, particularly those related to digestive enzymes and transport, would be impacted the most. While some of our hypotheses were confirmed, we were surprised by the rather limited transcriptional response to large changes in dietary p:c ratio. We were also surprised by the similarity in responses to diets that that varied from the intake target diet in opposite ways.

The numbers of DE genes observed across p:c ratio contrasts were relatively small. Less than 50 genes were up-regulated in the CB-42 treatment compared to the IT-42 diet and even fewer for the PB-42 treatment (though it must be acknowledged that the p-biased diet was more similar to the IT-42 diet than the c-biased diet; 6% difference in protein and carbohydrates for the CB-42 diet compared to 12% for the PB-42 diet). This was in stark contrast to impact of diet total macronutrient content, which showed the differential regulation of over 1,800 genes. When you consider the proportion of genes that were uniquely associated with each diet treatment, numbers are further reduced to only 3 up-regulated and 29 down-regulated genes unique to the CB-42 treatment and 4 up-regulated and 3 down-regulated genes unique to the PB-42 treatment (Figure 2). A greater number of down-versus up-regulated DE genes was observed in both the CB-42 and PB-42 diet treatments, which shows that feeding on a diet that differs from the intake target ratio has a suppressive effect on gene expression. This may indicate a strategy to limit physiological responses to nutritional imbalance in order to conserve resources. While data on the connection between nutritional state and the metabolic cost of generalized transcriptional responses is hard to find, the growth-rate hypothesis, which proposes a functional connection between dietary demands for phosphorus, endogenous concentrations of phosphorus-rich ribosomal RNA, and growth rates capabilities (Main et al., 1997; Sterner and Elser, 2002; Elser et al., 2003; Hessen et al., 2007), does provide a precedent for making connections between nutrition and physiological limitations. This notion is supported by the fact that many genes related to protein-intensive cellular components or processes, such as protein-containing complex, antioxidant activity, detoxification, reproduction, and immune system process, were down-regulated in the CB-42 treatment and were up-regulated in the PB-42 diet treatment (Table 2).

Despite the fact that the CB-42, PB-42, and IT-68 diets varied from the IT-42 diet in different ways, there were some surprising similarities in the transcriptional responses in these contrasts. Seven percent of up-regulated genes and 12% of down-regulated genes were uniquely expressed in the CB-42 diet, while only 18% of up-regulated genes and 3.5% of down-regulated genes were uniquely expressed in the PB-42 diet (Figure 2). No up-regulated DE genes were solely shared by these two contrasts and only 5 down-regulated DE genes were.

There was also considerable overlap in the function of these genes. Table 2 shows that the majority of GO terms associated with the DE genes from both ratio contrasts were shared. Interestingly, a high proportion of the DE genes from the p:c ratio contrasts were also shared by the total macronutrient contrast (IT-68 vs IT-42). Among up-regulated DE genes, 27.9% of those from the CB-42 vs IT-42 contrast and 54.4% of the PB-42 vs IT-42 contrast were also shared by the IT-68 vs IT-42 contrast. An even greater proportion of down-regulated DE genes were shared across all three contrasts; 31.5% of all CB-42 vs IT-42 DE genes and 88.2% of all PB-42 vs IT-42 DE genes. This is surprising given that all three diets differ from the IT-42 diet in completely different ways. These results suggest that similar mechanisms are utilized to deal with deviations away from the intake target, regardless of how a diet is different. These results also indicate that the physiological mechanisms evolved to mitigate nutritional variability are centered around a nutritional optimum (i.e. intake target), and further supports the importance of nutrient regulation in insects (Illius et al., 2002; Simpson et al., 2004; Simpson and Raubenheimer, 2012).

The impact of dietary total macronutrient content had a stronger impact on gene regulation than p:c ratio, resulting in the regulation of between 6.6 and 17.5 times more genes than the p:c contrasts (Fig. 2). These DE genes corresponded to a diversity of different GO terms, including many higher order processes. The strong transcriptional response to increased total macronutrient content was somewhat unexpected, given that the value of 42% is at the higher end of the range found in plant tissues (Deans et al., 2016, 2018; Lenhart et al., 2015) and not likely limiting, particularly if you consider that artificial diets are more easily digested than plant tissues (Lee at al., 2004; Clissold, 2007). Such a strong transcriptional response suggests that many physiological processes are routinely constrained by total macronutrient content and that larvae have evolved mechanisms to take advantage of high calorie foods when they are available. This is further supported by the fact that Deans et al. (2017) did not observe a reduction in consumption for larvae fed on the p39:c29 diet (comparable to our IT-68 diet), despite it being more concentrated.

Another surprising result was the limited regulation of metabolic genes, i.e. those involved in digestion, transport, and energy metabolism across diets. In terms of differential expression, few genes encoding digestive enzymes were found among the DE genes across contrasts (Figure 6), and the enrichment terms associated with DE genes in the p:c contrasts also identified few metabolic functions (Table 2). Figure 7a showed that the total number of protease transcripts did not vary across diets, despite an over 3-fold difference in dietary protein content. However, while there is some evidence that the differential expression of individual metabolic genes was not strong enough to indicate statistically significant differences in expression, the aggregate expression of genes within specific functional groups did show significant differences across diet treatments. For instance, less than four DE genes related to glycosidase activity were found across all contrasts (Figure 6), but the total expression level of all glycosidase transcripts did vary significantly between some diets (PB-42 and IT-68) (Figure 7a). Also, the GSEA, which analyzes aggregate expression across gene sets whether or not the genes have been found to be differentially-expressed, did uncover a significant correlation between several protease gene sets and the CB-42 (aminopeptidase), PB-42 (all protease gene sets), and IT-68 (trypsin, chymotrypsin, and carboxypeptidase gene sets) diets (Table 2). There was also a significant correlation between these three diets and the energy metabolism gene set. The IT-68 diet contained a greater total macronutrient concentration and thus, increased energy metabolism would be expected, however, increased expression of energy metabolism genes in the CB-42 and PB-42 diets may reflect an increase in energy expenditure associated with mitigating nutritional imbalances.

*H. zea* larvae are able to maintain larval performance across a broad range of dietary p:c ratios, exhibiting incredible nutritional plasticity (Deans et al., 2015, Deans et al., 2017). When presented with multiple diets that differ in p:c ratio, *H. zea* larvae feed selectively to ingest an optimal balance of dietary protein and carbohydrates (Waldbauer et al., 1984; Cohen et al., 1987; Deans et al., 2015); a response that has been observed in virtually all other insect species tested (Behmer, 2009; Simpson and Raubenheimer, 2012). Deans et al. (2017), however, showed that when limited to one diet, mass-specific consumption did not change across the same diets tested in this study. This produced significant differences in mass-specific protein and carbohydrate intake across diet treatments, proportional to protein and carbohydrate content in each diet (Table 3). Our data show that despite significant differences in protein and carbohydrate intake, there was little change in the cumulative expression of protease or glycosidase genes across diets (Figure 7a). A similar result was documented for trypsin activity in *H. zea* by Broadway and Duffey (1986). We did, however, find that when expression of protease and glycosidase genes was scaled to the amount of protein and carbohydrates ingested, there were significant differences in protease and glycosidase gene expression (Table 5).The carbohydrate-biased CB-42 treatment showed greater expression of protease transcripts per mg of protein ingested, and the protein-biased PB-42 treatment showed a greater number of glycosidase transcripts expressed per mg c ingested (Figure 7b). Therefore, maintaining consumption and consistent protease/glycosidase expression across different diets results in a disproportionately higher level of gene expression for enzymes pertaining to the more limiting macronutrient. Thisprovides an example of a compensatory mechanism that is mediated by maintaining consistent feeding behavior and constitutive expression of digestive enzymes, rather than by inducing changes in expression.

There is a wealth of data available on the relationship between host plant use and digestive enzyme activity in generalist lepidopterans (Patankar 2001, Kotkar 2009; Chougule 2005), as well as gene expression across host plant consumption. Most of these studies, however, focus on the effects of plant defensive compounds, including proteinase-inhibitors (Brioschi et al., 2007; Chikate 2013; Kuwar et al., 2015) and allelochemicals (Li et al., 2002; Pachet et al., 2010; Celorio-Mancera et al., 2012; Vogel et al., 2018) rather than nutrients. In fact, because differences in defensive compounds and nutrients between host plants are rarely quantified and reported, it is impossible to tease out the nutritional contribution to these transcriptional changes. Because we utilized artificial diets containing no plant components, we were able to explicitly measure the impact of specific dietary parameters.

Although we expected that the nutritional plasticity observed in *H. zea* larvae would be mediated by strong transcriptional changes in metabolic pathways, we saw a rather limited response in the number of diet-associated DE genes, as well as substantial similarities across responses to p:c ratio and total macronutrient content. Regulation of processes able to compensate for nutritional imbalances were not observed on a transcriptional level but were seen when the expression of digestive enzymes was corrected for macronutrient intake. It is unclear whether a strategy of fixed consumption and constitutive expression of digestive enzymes is common among insect generalists. Other polyphagous species, such as *S. exigua*, appear to show a stronger proteolytic response to increases in dietary protein (Broadyway and Duffey, 1986; Roy et al., 2016), but as mentioned, there are few studies that explicitly look only at dietary nutrients. Transcriptional studies that explore how other caterpillar species, particularly specialists, mediate nutritional imbalance will further uncover mechanisms associated with host plant adaptation and the evolution of monophagy/polyphagy. Additionally, plant allelochemicals play a significant, if not primary, role in host plant adaptation and must be incorporated into future studies to truly understand the evolution of feeding strategies and insect nutritional ecology. This is particularly true given that few, if any, insect herbivores feed on resources that don’t also contain some kind of allelochemical. Because metabolic responses to nutrients have evolved in the context of these other stressful plant compounds, it is possible that nutrient-mediated changes in transcription may only be triggered when these stressors are present and are undetectable under the rather unnatural control conditions tested in this study. Research suggest that the expression of detoxification enzymes, such as cytochrome P450s, esterases, glutathione S-transferases, UDP-glucuronosyltransferases, ATP-binding cassette transporters, etc., are responsive to differences in host plants (Li et al., 2002; Pachet et al., 2010; Celorio-Mancera et al., 2012; Koenig et al., 2015; Vogel et al., 2018) but interactions between nutritional state and responses to allelochemicals are still ambiguous (Deans et al., 2016, 2017). Ultimately, the interplay between minimizing consumption of allelochemicals, mounting an effective physiological response to allelochemicals, and allocating resources to detoxification processes must be more thoroughly studied.

## ACKNOWLEDGEMENTS

This work was supported in part by the C. Everette Sayler Fellowship, the Charles R. Parencia Endowment through the Entomology Department at Texas A&M University, and a Biotechnology Risk Assessment Grant Program competitive grant no. 2015-33522-24099 from the U.S. Department of Agriculture (awarded to GAS, STB, and MP-C)

---

**SI Fig. 1.**
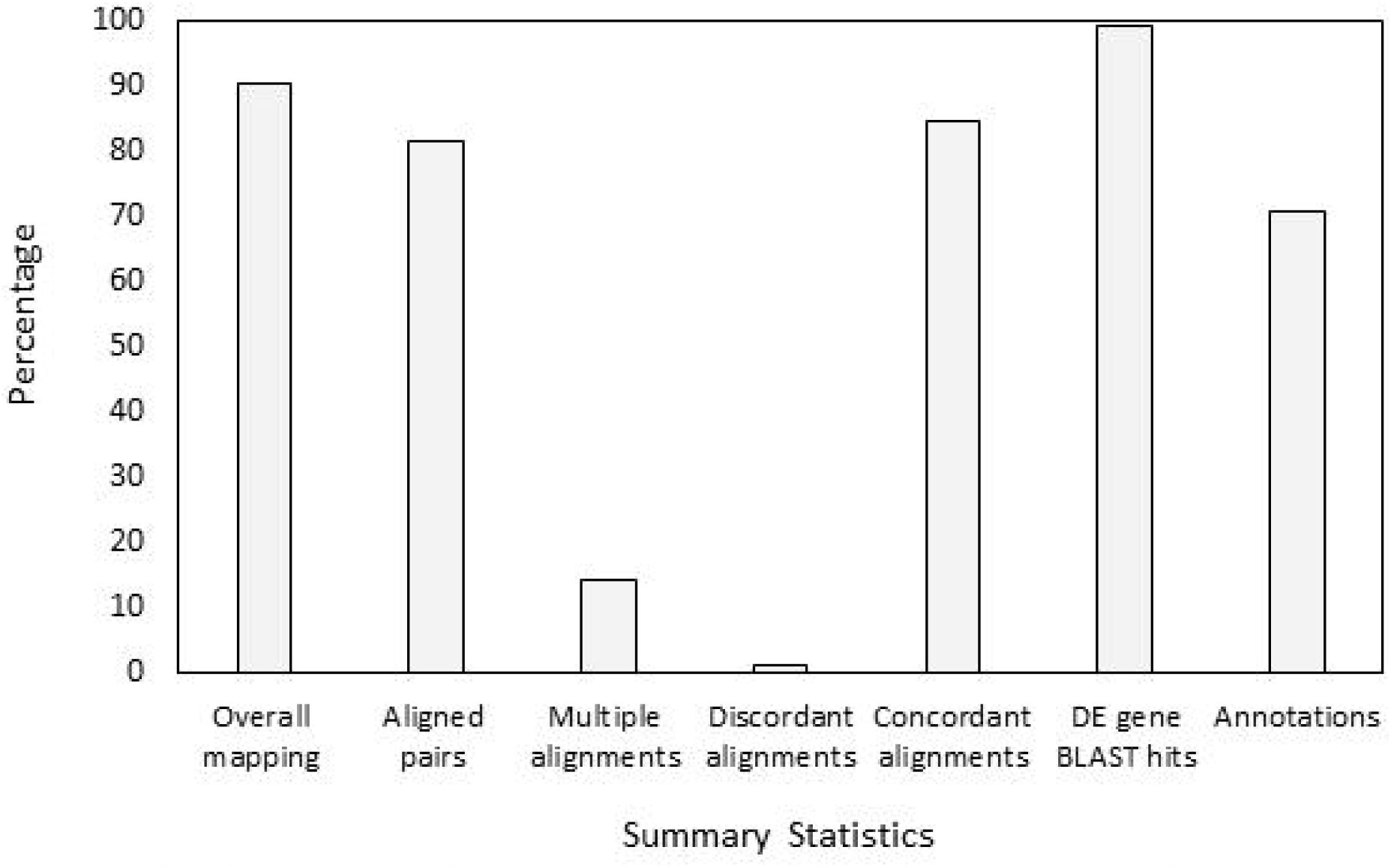
Summary statistics. Graph showing a summary of the alignment, mapping, and annotation statistics.

**SI Fig. 2.**
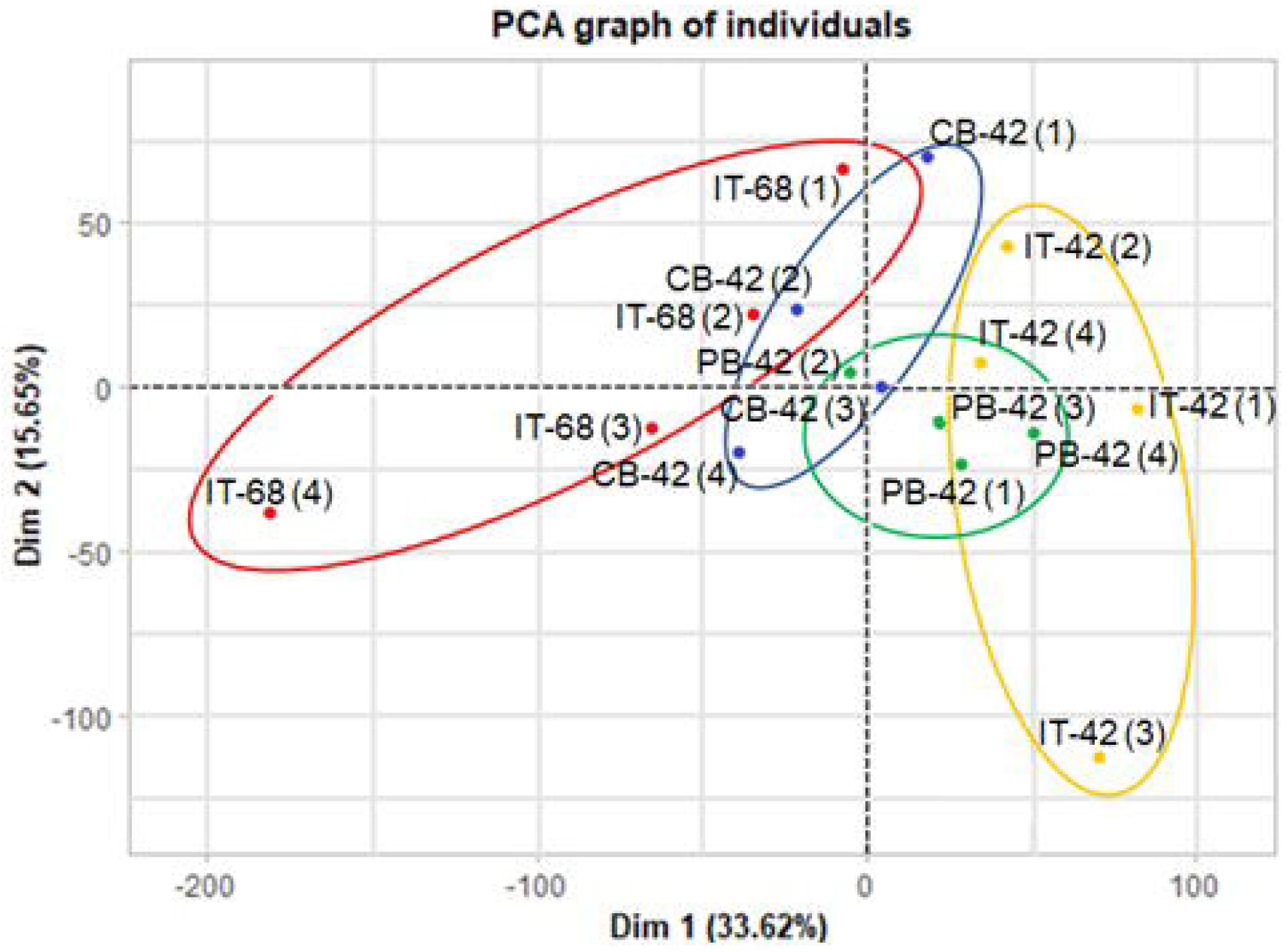
PCA plot of all replicates. This plot shows the replicate clusters for each diet treatment: IT-42 (yellow), CB-42 (blue), PB-42 (green), and IT-68 (red).

